# Host control and species interactions jointly determine microbiome community structure

**DOI:** 10.1101/2022.03.03.482885

**Authors:** Eeman Abbasi, Erol Akçay

## Abstract

The host microbiome can be considered an ecological community of microbes present inside a complex and dynamic host environment. The host is under selective pressure to ensure that its microbiome remains beneficial. The host can impose a range of ecological filters including the immune response that can influence the assembly and composition of the microbial community. How the host immune response interacts with the within-microbiome community dynamics to affect the assembly of the microbiome has been largely unexplored. We present here a mathematical framework to elucidate the role of host immune response and its interaction with the balance of ecological interactions types within the microbiome community. We find that highly mutualistic microbial communities characteristic of high community density are most susceptible to changes in immune control and become invasion prone as host immune control strength is increased. Whereas highly competitive communities remain relatively stable in resisting invasion to changing host immune control. Our model reveals that the host immune control can interact in unexpected ways with a microbial community depending on the prevalent ecological interactions types for that community. We stress the need to incorporate the role of host-control mechanisms to better understand microbiome community assembly and stability.

## 2 Introduction

One of the fundamental questions concerning ecology is understanding the processes influencing species diversity in communities. The field of theoretical ecology has mainly addressed this question using a resource-centric approach introduced by Lotka and Volterra (Kopf and Lotka, 1925, Volterra, 1926). Later, MacArthur and his colleagues using the Lotka-Volterra framework formulated a model for resource competition where species with overlapping resource requirements competitively exclude the other based on each species carrying capacity and competition coefficient (Macarthur and Levins, 1967, MacArthur, 1970). More recent work, building on May’s random matrix approach (May, 1972, 1973), considered the dynamics of complex communities with varying prevalence of interaction type such as mutualism, exploitations, and competition (Melián et al., 2009, Mougi and Kondoh, 2012, Coyte et al., 2015, Qian and Akçay, 2020). These studies show that the balance of interaction type influence the community dynamics, species richness and the internal and external stability of the community.

Microbial communities can be studied using this community ecology framework (Costello et al., 2012, Gilbert and Lynch, 2019), but with a twist. The community dynamics of the host microbiome are governed not just by how species within the microbiome interact with one another, but also by the interaction with the host organism. The species interactions in the microbiome can range from resource mediated competition to cooperative and exploitative interactions (Foster and Bell, 2012, Gralka et al., 2020). Many microbial species are auxotrophic - lack essential genes to produce metabolites including vitamins and proteins for cell growth and survival (Zengler and Zaramela, 2018). These species have to rely on other microbial species to fulfil their nutritional requirements and also engage in the successive digestion of large complex molecules provided by the host (Levy and Borenstein, 2013, Zengler and Zaramela, 2018, Gralka et al., 2020). The host can mediate the community structure and assembly of such communities through the supply of these resources. The host can also affect microbiome assembly and stability through its immune response which can be specific or general (Hooper et al., 2012, Stagaman et al., 2017).

Microbiome dynamics are under the joint control by both their hosts and within-microbiome ecological interactions. The host in particular is under selective pressure to support a microbial community that is compatible with the host, which should select for mechanisms that regulate community assembly, diversity, and stability (Foster et al., 2017). In essence the host-associated microbiome can be considered as an ecosystem held on a leash (Foster et al., 2017), where the leash signifies that the host will attempt to keep microbial communities within certain bounds. The host can tighten or loosen its leash on the microbiome that can consequently affect the microbial species that are able to enter the host and eventually colonize within a region such as the gut.

Here, we present a model of the microbiome as an ecosystem on a leash using a generalized Lotka-Volterra framework (Bunin, 2017), with type-II functional responses. The Lotka-Volterra framework has been applied to infer and predict microbial population dynamics (Foster and Bell, 2012, Stein et al., 2013). However, this framework has mainly been applied to study ecological interactions between microbial species without much consideration of the host control over the microbial community. Specifically, we consider immune responses as the leash that the host holds to control the assembly of microbial communities.

How host immune system controls microbiota can be complex and variable across species, and is still not fully understood. Inside the gastrointestinal (GI) tract of mammals, the host’s epithelial barrier and the mucus serve as the first layer of physical separation between the microbiota and the host tissues (Hooper et al., 2012, Belkaid and Hand, 2014, Stagaman et al., 2017). In the event that microbes translocate beyond the epithelium barrier, the host immune system recognizes microbial-associated molecular structures using toll-like receptors and targets these microbes by releasing inflammatory cytokines, macrophages, and immunoglobulin A (IgA) (Belkaid and Hand, 2014). Hosts can also vary the environmental pH (Ratzke and Gore, 2018) and oxygen levels (Byndloss et al., 2018), restricting the growth and persistence of specific microbial species. These physiological and environmental components are instances of host control that can impose negative selection on microbes. However, how this control varies in response to a beneficial symbiont or a pathogen to the host is unclear. The distinction between a pathogen or a beneficial symbiont to the host can be challenging for the host immune response as pathogenicity is context dependent, and there are known instances where pathogens are able to proliferate even in the presence of an active immune response (Rivera-Chávez and Bäumler, 2015). Moreover, host control over the specific microbial community members is less evident from human fecal microbiota analysis (Tap et al., 2009). Invertebrate hosts that lack an adaptive immune response are more likely to rely on non-specific host immune responses. Invertebrates such as corals have innate immune responses that can sense microbial molecular patterns and release antimicrobial peptides which regulate microbial loads in a non-specific manner (Palmer et al., 2011, Palmer, 2018). These observations suggest that host imposing a generalized, non-specific host immune response is likely to be an important mechanism of regulating microbial community abundance in host-associated microbial ecosystems.

Here, we consider host control at the level of the whole microbiome, where the host regulates the microbiome density through resource provisioning and the immune response. Microbial community density is a fundamental ecosystem property that can have implications on the host health. How the microbial community responds to host control relies not just on the type of microbial species present, but also on the abundance of these species. Contijoch et al. (2019) observed a natural variation in the gut microbiome density across a range of mammalian hosts, with each host supporting a specific microbial carrying capacity. There is also variation observed in microbiome density observed across host organs, where in humans the bacterial colony forming units increase from stomach to small and large intestine (DiBaise et al., 2006), indicating differential host control at play at the organ level. Contijoch et al. (2019) report a positive correlation between immune cell population and the microbiome density in specific pathogen free (SPF) mice treated with varying density depleting antibiotics. Therefore it is probable that changes in host control can alter the microbiome density, that can consequently affect the host health.

In this paper, we present a mathematical framework to model host-associated microbial community dynamics. Understanding microbial community dynamics is essential as the microbiome harbors a vast diversity of functionally important microorganisms ranging from bacteria, archaea, and fungi (Huttenhower et al., 2012). These microorganisms play a crucial role in providing host protection against pathogens, aid in immune maturation and metabolism (Lozupone et al., 2012). Changes to the stability and composition of the microbiome composition is associated with pathologies such as inflammatory bowel disease, diabetes (Zaneveld et al., 2017, Lozupone et al., 2012) and more recently cancer (Helmink et al., 2019). Existing genomics sequencing of microbial community provide limited understanding of the forces at play that shape the microbiome. Furthermore, existing experimental and theoretical approaches to study microbial community dynamics are unable to provide an understanding of the implications of the interaction between the host control and the balance of ecological interaction types. To address this gap, we apply and extend the framework by Qian and Akçay (2020) to model a microbial community driven by both within-species ecological interactions and host control. The host immune response can serve as an active modulator of the microbiome density that can have an affect on microbial community composition and assembly. To understand the interplay between host control and the assembly of the microbiome, we integrate global density dependence mediated by host immunity and ask how it interacts with the balance of species interactions within the community.

## 3 Methods

We model the host control and the sequential assembly of the microbial community extending the model by Qian and Akçay (2020). We simulate communities comprised of *S* microbial species. We initialize the community with *S*_*o*_ species, and the community grows in size through successive invasions, and can also decrease in size as extinctions occur. Microbial species can engage in pairwise interactions with other members in the community. We incorporate all possible pairwise combinations of ecological interactions; proportion of mutualism (*P*_*m*_), exploitation (*P*_*e*_), and competition (*P*_*c*_) when constructing the interaction matrix *A*. We vary the proportion of all ecological interactions in intervals of 0.1, with each microbial community having a unique *P*_*m*_, *P*_*e*_ and *P*_*c*_. All communities are exposed to some degree of competition (*P*_*c*_ ≠ 0). We ensure that *P*_*m*_, *P*_*e*_ and *P*_*c*_ chosen for each community all sum to one.

The mathematical equation for population dynamics of microbial species is given below:

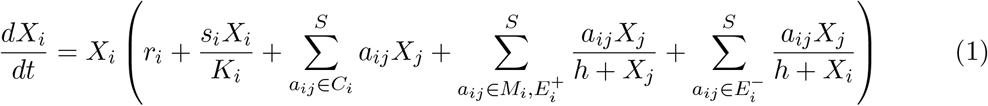

where

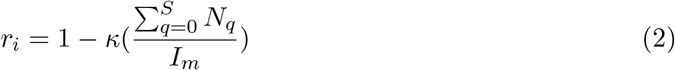

*X*_*i*_ represents the population of species *i*, and *r*_*i*_ represents the intrinsic growth rate. We modify the growth rate *r*_*i*_ (Equation2) of all microbial species as an instance of immune-mediated host control over the microbial community size. The host immune control is synonymous to introducing global density dependence which ensures that microbial community abundance dose not exceed the host specific microbial carrying capacity (*I*_*m*_). We also introduce *κ*, which represents the reduction in all microbe growth rates with the total community abundance and can be thought of as the inverse of a community-wide carrying capacity. *N* represents the species population. When simulating a mammalian gut community we choose *I*_*m*_= 10000, as the bacterial CFUs per milliliter for the gut are estimated at this magnitude (Miller and Baumler, 2021). Changes to *I*_*m*_ still preserves the qualitative results of our model.

The coefficients *a*_*ij*_ characterize interactions between species *i* and *j*. For mutualistic and antagonistic interactions, we use Type II functional responses, with *h* as the half-saturation constant. We denote by *s*_*i*_ the self-regulation term and *K*_*i*_ the species-specific carrying capacity. *C*_*i*_ is the set of interactions between the *i* and its competitors, *M*_*i*_ is the set of interactions between *i* and its mutualistic partners, *E*^+^ represents the set of interaction between *i* and the species it exploits, whereas *E*^*−*^ refers to the set of interactions between species *i* and species that exploits it. These equations correspond to a generalized Lotka-Volterra model (Bunin, 2017), but with the crucial difference of non-linear functional responses. The saturating functional responses, coupled with the sequential assembly of the communities (which selects for feasible communities) result in almost all our assembled communities being asymptotically stable (Roberts, 1974, Stone, 2018, Qian and Akçay, 2020, see also SI.10, SI.11).

### 3.0.1 Community simulation

The initial population of each species is set to a constant *x*_*o*_ multiplied by a random number from a uniform distribution N(0,1). For all species we set a constant carrying capacity, *K* = 100, and *r* = 1 at the start for the microbial community. We initialize the interaction matrix *A*_*o*_ with dimensions *S*_*o*_ by *S*_*o*_. The diagonal values of the interaction matrix are set to *s* = *−*1. The interaction between each species pair i,j where *i* ≠ *j* is determined if the random probability drawn from normal distribution N(0,1) is less than or equal to the connectivity term, *c*. If no interaction exists then *a*_*ij*_ and *a*_*ji*_ are assigned a zero value. For interacting species, species *i* and *j* can compete with each other with *P*_*c*_. These species are assigned *a*_*ij*_ and *a*_*ji*_ values drawn from a negative half normal distribution -N(0, *σ*/*K*). The species that have a mutualistic interaction with probability *P*_*m*_, are assigned *a*_*ij*_ and *a*_*ji*_ values drawn form a half normal distribution N(0, *σ*=0.5). Species can also engage in an exploitation interaction with a probability *P*_*e*_, in this scenario, we randomly assign species *i* and *j* to be either exploited or to exploit the other species. In the case if species *i* is being exploited then *a*_*ij*_ has a value drawn from an -N(0, *σ*) and *j* being the exploiter, has *a*_*ji*_ from N(0, *σ*).

After the initial setup of the interaction matrix *A*_*o*_, we then simulate the population dynamics of all *S*_*o*_ species until the population has reached equilibrium. We integrate the microbial species population dynamics using the Dopri5 integrator. We determine if the population has reached equilibrium: i) if at a single time step the population fluctuates less than the population change threshold *δ*, then the population has reached equilibrium, (ii) if the population has not reached the equilibrium and the time limit, *t*_1_ has been reached then we use the current population sizes of the species in the community as the equilibrium values. We introduce a new species into the community once the equilibrium is reached. At each equilibrium before introducing a new species we draw a small population of the potential invader and consider it along with the abundance of the resident microbial species to update We draw new interaction coefficients for the potential invader, similar to how we draw during the initialization of the interaction matrix *A*_*o*_. An invader can only colonize if the growth rate of the invader is positive when a small population of the invader is added into the community, accounting for the invader’s interactions with the existing resident species. If invasion fails, then we redraw the interaction coefficients of the invader until it is able to invade. We impose a failed invasion threshold limit, *β*= 1000. If *β* is reached then we end the simulation and deem that equilibrium externally stable-uninvadable. If a successful invasion has occurred, we update the interaction matrix *A*, to now have dimensions of *S*_0_ + 1 by *S*_0_ + The rows and columns of *A* are updated such that to conserve the given *P*_*m*_, *P*_*e*_ and *P*_*c*_ of the community. We ensure that the interactions between the resident species do not change during invasions. We simulate population dynamics until next equilibrium, after which we introduce a new species. We also determine the community’s competitive barrier to invasion by summing up the effect of all the competitive interactions(*a*) between the invader (*i*) and the resident species (*j*) scaled by the resident species abundances (*X*) in the community:

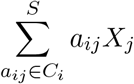

Species also experience a decrease in their abundance and hence can go extinct. If the species abundance is lower than the extinction threshold *ϵ* then the species is deemed extinct and is removed from the community. Throughout the simulation we keep track of the number of species that went extinct, the number of invasions and the microbial species richness observed for each microbial community. We then use these values to determine the probability of extinction (number of extinct species over the number of species observed at that point of community history) and invasion (number of invasions over all invasion attempts) for the community.

Species richness of the microbial communities can converge, and to assess the presence of a steady state of species richness we use Dickey-Fuller test, as implemented by (Qian and Akçay, 2020). We analyze the most recent 1000 equilibria and determine if the P value is less than the threshold *p* value, that indicates that the species richness has entered the steady state. Once a community enters the steady state we run the community for *w*_2_ more equilibria before we end the simulation. We terminate simulations with communities that never enter the steady state if the community reaches (i) limit on number of species (*n*), (ii)limit on the number of equilibria (*τ*), or (iii) *t*_1_.

We run five simulations for each microbial community, so we end up in total with 55×5 = 275 microbial communities. The results presented in the paper are an average of 5 simulation runs. Table 1 represents the parameters used in the model.

**Table 1:** Fixed variables in the model and their definitions.

| Model Fixed Parameters |  |
| --- | --- |
| Variable | Definition |
| $s_i = -1$ | self regulation term |
| $h = 100$ | half-saturation constant of type II functional response |
| $c = 0.5$ | connectivity term of the microbial species |
| $K_i = 100$ | species-specific carrying capacity |
| $\epsilon = 0.001$ | failed extinction threshold |
| $\beta = 1000$ | failed invasion threshold |
| $\tau = 5000$ | equilibria limit |
| $p = 0.01$ | p value threshold |
| $I_m = 10000$ | host immune supported carrying capacity |
| $n = 750$ | species number limit |
| $w_1 = 1000$ | number of equilibria analyzed for the Dickey fuller test |
| $w_2 = 500$ | number of equilibria simulated after steady state is reached |
| $\delta = 0.001$ | population change threshold |
| $t_1 = 10^9$ | time limit for simulation |
| $\sigma = 0.5$ | half-normal distribution scale parameter |

## 4 Results

We simulate the sequential bottom-up assembly of microbial communities with varying fractions of all possible interaction types that include competition (at fraction *P*_*c*_ of all species interactions), mutualism (at fraction *P*_*m*_), and exploitation (at fraction *P*_*e*_), as in Qian and Akçay (2020). We initialize a microbial community with a *S*_0_ number of species, and let the community grow in size through successive invasions. For each invasion event, we draw interaction coefficients for the invader from a random distribution where the probability of an interaction being of a given type is given by the fractions *P*_*c*_, *P*_*m*_, and *P*_*e*_. If the invader can increase from rare, we numerically evaluate the new community dynamics until it reaches an equilibrium and remove species that fall below an extinction threshold at this equilibrium. If the invader cannot increase from rare, we draw a new invader species. In this way, community size can grow or shrink through invasions and extinctions.

To simulate community dynamics, we integrate microbial within-species interactions and the host immune control in a Lotka-Voltera framework, where microbial species densities are regulated by a combination of intra-specific density dependence, between species interactions, and host immune control. We model the host immune control as responding to the total microbial community size. Specifically, we assume that all species’ growth rates decline as a function of the overall community abundance, amounting to a global negative density-dependence term. The strength of host control strength is given by *κ*, which represents the reduction in all microbe growth rates with the total community abundance and can be thought of as the inverse of a community-wide carrying capacity. Higher *κ* corresponds to stronger host control.

In absence of host control, we recover the results of Qian and Akçay (2020). The microbial community species richness is governed by a balance of interaction types, where more mutualistic and less competitive microbial communities (high *P*_*m*_ and low *P*_*c*_) harbor the highest number of microbial species (Figure 1a, at *κ* = 0). In our model, microbial species richness emerges as a balance between the number of invasions and extinctions of species in a community. Highly mutualistic microbial communities are colonization resistant with a low probability of invasion (Figure 1b, at *κ* = 0). This is because communities with a high proportion of mutualistic/cross-feeding interactions are able to achieve a high community density. Mutualistic interactions positively contribute to the growth rate of the interacting species, building up resident species equilibrium population sizes in the community (Figure SI.2b, at *κ* = 0). This also explains why highly mutualistic communities have a shallow rank abundance curve slope when compared to the steep slope which is indicative of low species evenness observed for microbial communities with high *P*_*c*_ (Figure 2b). Communities with high *P*_*m*_ and low *P*_*c*_ are still able to achieve a high competitive barrier to invasion due to the high equilibrium population size of resident species (Figure 1c, at *κ* = 0) buffering the resident community from getting out competed by incoming microbial species (Figure SI.2a, at *κ* = 0).

**Figure 1:**
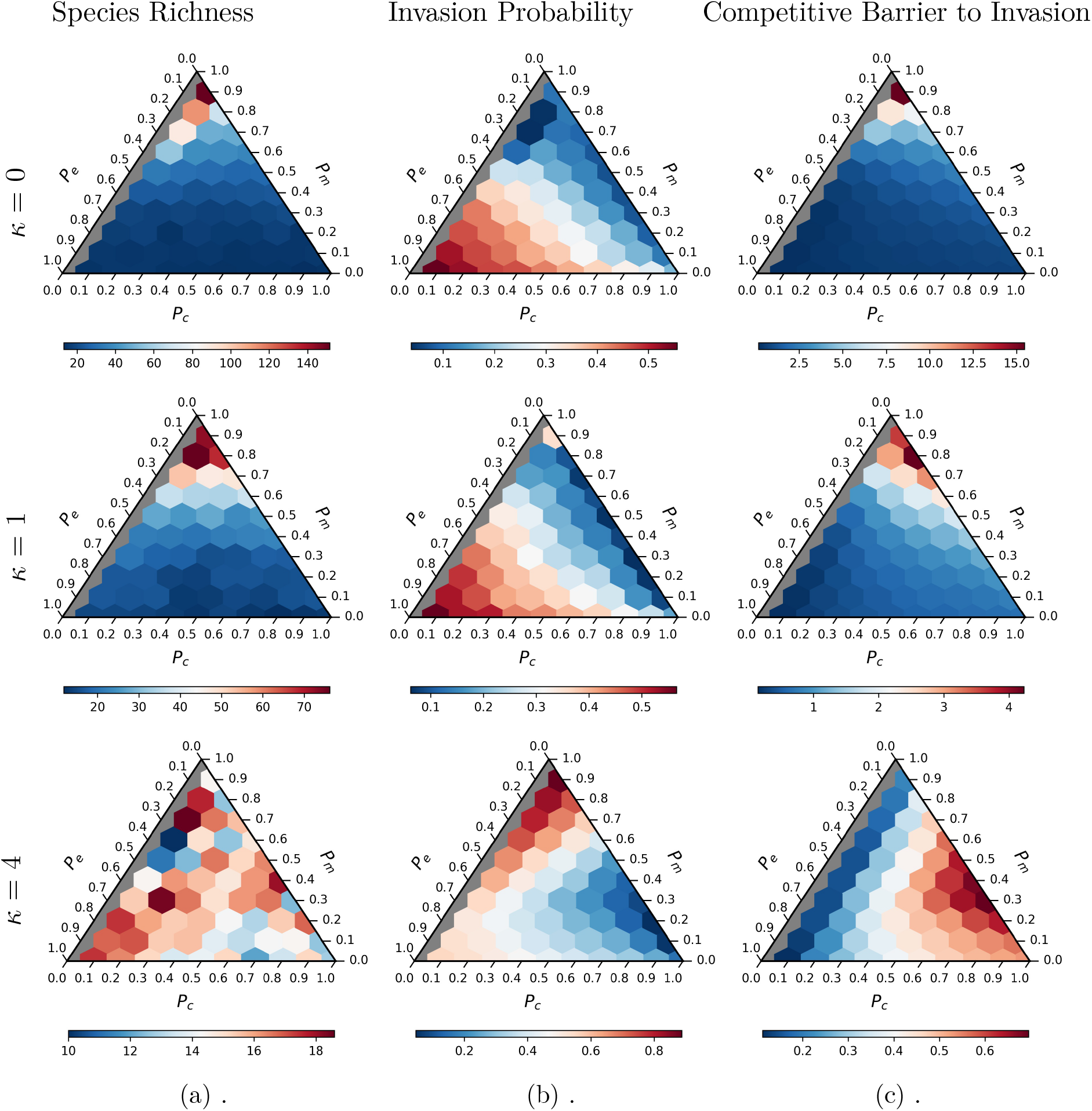
Ternary plots representing the host control on (a) species richness (at steady state), (b) invasion probability at steady state, and (c) competitive barrier to invasion for all possible communities with different balance of interaction types. In each ternary plot, the location of each hexagon corresponds to a particular balance of interaction types, given by probabilities *P*_*c*_, *P*_*m*_ and *P*_*e*_ for competitive, mutualistic, and exploitative interactions, respectively. Note that scale of each plot shown above has a different range. Each row demonstrates an increasing host control strength (*κ*) for each community metric, from no host control (*κ* = 0) to high host control strength (*κ* = 4).

**Figure 2:**
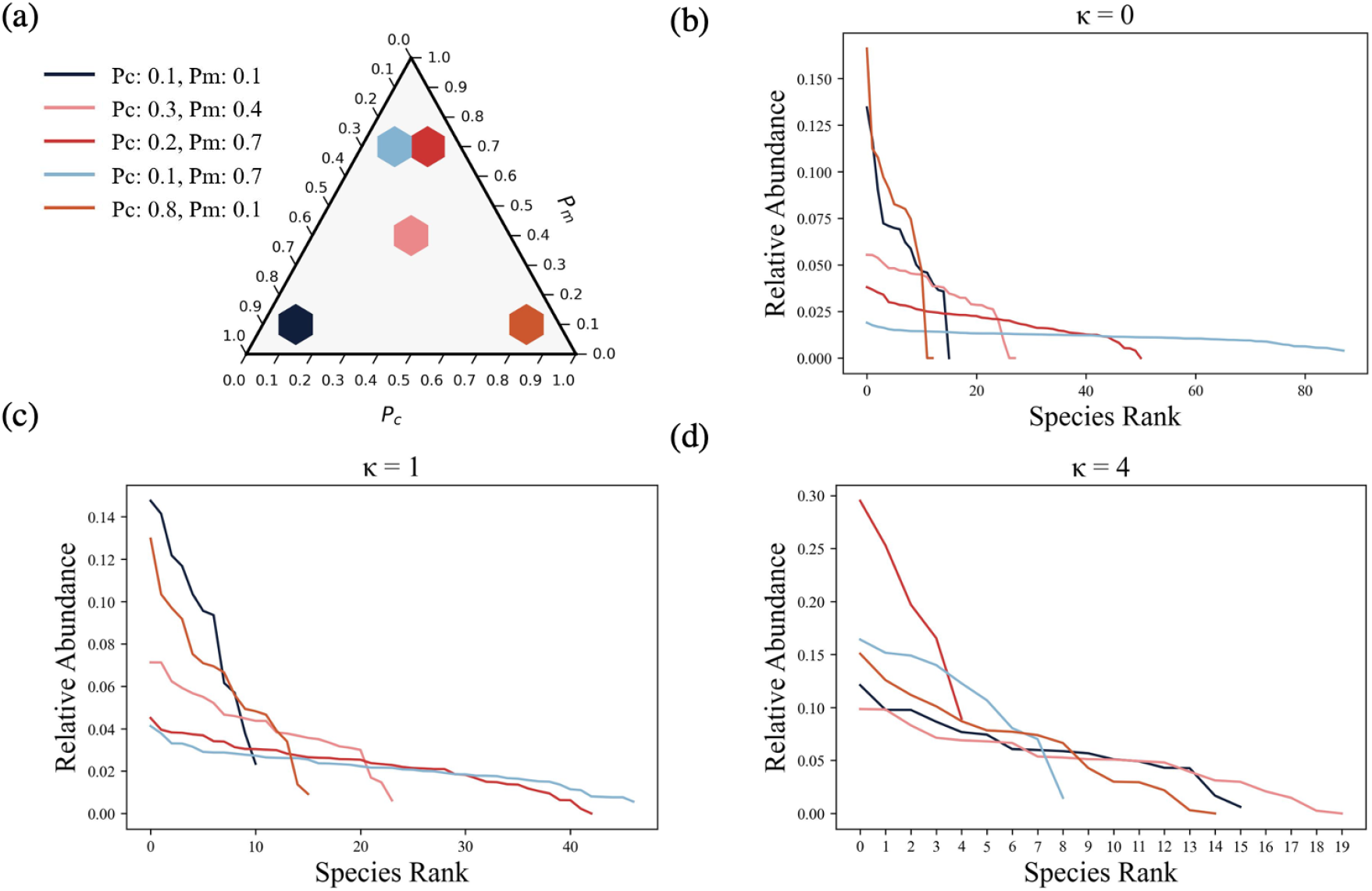
(a)Represents a ternary plot with selected microbial communities. The color of each selected community in the ternary plot corresponds to line plots in parts b-d. The line plots represent species rank abundance curves across a range of *κ* values. Note that scale of each plot shown above has a different range.

Introducing host immune response has non-trivial effects on community dynamics. As we introduce host control (*κ >* 0), we observe that microbial communities’ species richness generally decrease. This decrease is most marked for communities with high *P*_*m*_ (Figure 1a, at *κ >* 0), whereas species richness for communities with high *P*_*c*_ remains relatively unaffected to changes in host control (Figure SI.1b). At very high host control strength, species richness of all communities converges to species richness observed for highly competitive microbial communities (Figure SI.1b). This suggests that elevated host control overrides the effects of the balance of interaction types in determining microbial species richness. This also results in the similar species evenness observed across microbial communities regardless of the interaction type when *κ* is further increased (Figure 2c-d). In our community simulations we have both strong global density dependence through host immune response and highly mutualistic interactions within a community at the same time. These two forces pull in opposite directions: one working to reduce community size and the other increase community size. As a result, we observe communities with a distinct cyclical pattern of species richness as new species are introduced into the community (Figure 3 at *κ* = 1), and the frequency of the cyclical pattern increases as we further increase the host immune response strength (Figure SI.4). The pattern emerges as an invasion event causes the extinction of the resident community (Figure SI.2a, at *κ >* 0), lowering the community abundance until the community grows back again and becomes increasingly susceptible to the density-regulating effects of the host immune response. We do not observe this pattern with competition alone (Figure 3). Thus, mathematically the incorporation of the host immune term in our model framework is not equivalent to further increasing the competition among the species.

**Figure 3:**
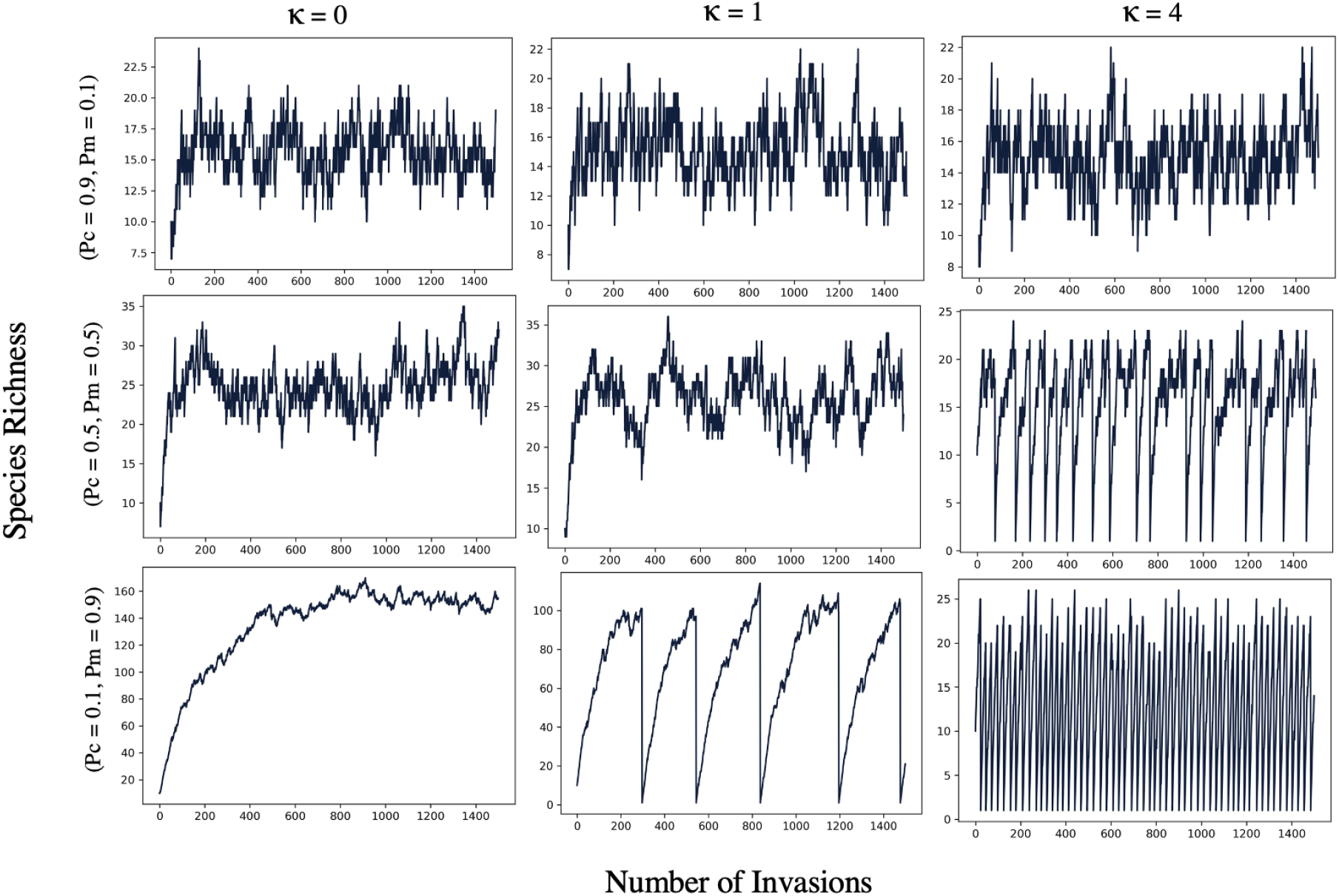
A series of line plots representing species richness for a microbial community as invaders are introduced into the community. Each row of the line plot represents a microbial community with a distinct *P*_*c*_, *P*_*m*_, and *P*_*e*_ across a range of *κ* values. The cooperation (*P*_*m*_) term increases and the competition (*P*_*c*_) decreases across rows of the line plots. Note that y-axis scale of each line plot shown above has a different range.

Host control pushes the intrinsic growth of all microbes to negative levels when the total microbial community abundance is greater than the host target carrying capacity. This reduces overall community sizes relative to no host control, and has the largest effect on the stability of highly mutualistic communities that rely on large community sizes to provide competitive exclusion to invading species (Figure SI.1d at *κ ≥* 1). The high total abundance that communities with high *P*_*m*_ can reach without host control triggers the host immune response to lower the growth rate of all microbial species in the community (Figure SI.2c at *κ ≥* 1). As a result the community population size decreases, consequently lowering the community’s competitive barrier to invasion (Figure SI.2b and Figure 1c, at *κ ≥* 1). These communities become more vulnerable to extinction, and observe a decrease in species persistence (SI Figure SI.2d at *κ ≥* 1) as the probability of invasion for the community increases with an increase in host control strength (Figure 1b at *κ ≥* 1, Figure SI.1c). In contrast, host control does not have much of an effect on microbial communities with high *P*_*c*_ (Figure SI.1b-d). Such communities are already subjected to high levels of competition and a lower growth rate such that introducing host control which is analogous to incorporating apparent competition in the community further adds/intensifies the competition in a community that is already highly competitive. Highly competitive microbial communities do not reach elevated levels of microbial community abundance to begin with (Figure SI.2b), hence these communities do not trigger the host immune control response. To check for potential history-dependent effects, we also considered community assembly with time-varying host control strength. We found that past host control strength has minimal role in determining how the community responds to a change in *κ* once the community has reached a steady state with the new host control strength (Figure SI.6 - SI.9).

## 5 Discussion

One of the main findings of our model is that a strong, generalized host immune control weakens the influence of the balance of interaction types on the community level properties of the microbiome. The host control can alter intrinsic dynamics of the microbiome differentially based on the prevalent ecological interaction types in the community. Microbial communities with high levels of mutualism appear most susceptible to changes in the host immune control and become increasingly susceptible to invasion. The high density of resident species characteristic of mutualistic communities, triggers the host immune control to decrease the overall community density which consequently lowers the community’s competitive barrier to invasion. In contrast, competitive communities whose species density is already set lower by within-microbiome competition remain stable to changes in the host immune control. Machado et al. (2021) present empirical evidence for the prediction that microbial communities that cooperate harbor a high community density compared to competitive microbial communities that rely more on the availability of externally supplied resources. Hence both species interactions and host control can collectively determine the composition and external stability in microbial communities. Our results suggest that immune control is more relevant as a host control mechanism to regulate microbial communities that are engaged in more cooperative or cross-feeding interactions.

Our model also indicates that hosts can achieve microbiome stability against invasion of new species (such as pathogens) either by hosting competitive microbial communities, irrespective of host immune control strength, or by hosting highly mutualistic microbial community with minimal immune control. This poses a trade-off between the degree of host control versus colonization resistance for a mutualistic microbial communities. It is important to note that a colonization resistant microbial community may not necessarily be compatible with the host nutritionally and immunologically. Both host and the microbiome are subjected to distinct selection pressures, where microbial species despite providing benefit to the host can fail to persist if they are unable withstand the interspecies competition (Foster et al., 2017). Therefore a colonization resistant microbial community may not necessarily be beneficial to the host. Colonization resistance then comes at a price to the host, where high competitive barrier to invasion results in the host being unable to control the associated microbial community. Conversely, elevated host control over the microbial community weakens the competitive barrier to invasion for highly mutualistic microbial communities, lowering the resident microbial species richness and persistence. Hence a symbiotic relationship between the host and the microbiome for highly mutualistic microbial communities relies on the balance between the host immune control and the colonization resistance.

Changes to the balance between the colonization resistance and the host control over a microbial community can have implications for host health and the associated microbiome. Host control can be impaired by changes to the homeostatic immune state of the host (Miller and Baumler, 2021). Immunodeficiency – a defect in the immune control over the microbiome – can trigger dysbiosis of the microbial community. Immunodeficiency can result in the expansion of the microbial community in the small intestine more than what is observed at the healthy host state, resulting in hosts suffering from small intestine bacterial overgrowth (SIBO) disorder (Zaidel and Lin, 2003). The overgrowth of microbial population interferes with the nutrient absorption in the small intestine as the microbial community is in competition with the host for the nutritional resources. The microbial community with increased energy demands readily consumes carbohydrates, and lowers the availability and absorption of other nutrients such as fats, fat-soluble vitamins and proteins causing the host to suffer from malnutrition (Zaidel and Lin, 2003, Siniewicz-Luzenczyk et al., 2015, Miller and Baumler, 2021). The weakened host immune control as evident in the SIBO metabolic disorder might cause the host to be no longer able to restrict the microbial community density to be under the host-supported carrying capacity. A reduction in the background competition imposed by the host immune control results in an increase in the growth of opportunistic microbial species specifically favorable for those that are engaged in cooperative interactions with other members of the community.

Conversely, an overt inflammatory immune state indicative of an elevated host control results in a decreased microbiome density observed in hosts with an inflammatory bowel disease (IBD) (Kiely et al., 2018). Contijoch et al. (2019) found a decrease in absolute abundance of firmicutes, actinobacteria and bacteroidetes with depleted microbiota density in hosts with IBD, whereas proteobacteria sustained a constant density in individuals with IBD. This suggests that some species may be more resilient to changes in host immune control. In line with our findings, proteobacteria phylum is known to be less abundant, which may cause them to trigger less strong immune responses. Further proteobacteria were also more functionally variable at a gene level compared to other phyla (Bradley and Pollard, 2017) which suggests that these species may share fewer metabolic co-dependencies with other species and engage in more competitive rather than cooperative interactions. These patterns need to be experimentally validated to further our understanding of the host-mediated control on microbiota density on the community composition and assembly of the microbial community.

In addition to host immunity, the amount and type of resources supplied by the host can influence microbial community composition. (Saffouri et al., 2019) found that healthy individuals with no symptoms can also exhibit SIBO, which correlated with high-fiber diets rich in complex carbohydrates. Complex carbohydrates might be a resource with higher opportunities for metabolic interdependencies (Smith et al., 2019) and thus positive interactions in the microbiome. Thus, it is possible that a diet rich in complex carbohydrates can shift the balance of interactions from being more competitive to more mutualistic. In that case, our model predicts an increase in both microbiome density and diversity (Fig SI.1b) even in the absence of a change in host control. This is consistent with the finding that gut-derived microbial communities provided structurally complex carbohydrates maintain higher species richness in lab culture (Yao et al., 2020). Our model suggests that such an increase in mutualistic interactions can either reduce the external stability of the microbiome (increase invasion probabilities) or leave it unchanged, depending on the strength of host control (FigSI.1c). This points to the need for more work on how metabolic relationships in the microbiome change with host-provided resources and how these interact with immune mechanisms.

In our model framework we incorporate host control as a global density dependence term. Where host modulates the entire microbial community, indiscriminately. Other mechanisms of host control include host resource availability which can actively modify the landscape of within-species interactions within the microbial community. Resource-based host control may not just serve as a global ecosystem modulator, but may also impose positive and negative selection on specific microbial species in the residing microbial community. Moreover, the host control can also be modified by the microbial community itself. The microbiome can engineer its micro environment by producing metabolites, which in turn can influence the biotic interactions in the microbial community (Gralka et al., 2020, Marsland et al., 2020, Kurkjian et al., 2021). Microbial community members can also influence host immune homeostasis, as specific microbial species can induce host immune responses (Atarashi et al., 2013, Ivanov et al., 2009). Microbiome-induced host immune response can consequently affect the strength of host control over the microbial community. The effect of these myriad ways of host control needs to be theoretically and experimentally investigated to truly uncover the effect of host-microbiome feedback on the host control over the microbial community.

In conclusion, we present a community dynamics framework, incorporating the role of host immune control in regulating microbial community size. We show that host immune control can interact with the within-microbiome species interactions in unexpected ways influencing the microbial community composition and stability. Host immune control may serve as a more relevant host control instance to keep the size of highly mutualistic microbial communities in check. We stress that understanding microbial community assembly and stability is incomplete without considering the role of host-control mechanisms on the microbial community.

## 6 Code availability

The model of microbial community assembly was implemented in Python. The simulation code is available at github: https://github.com/erolakcay/MicrobiomeCommunityAssembly

## 7 Acknowledgements

We thank Katie Barott for her helpful comments on the manuscript.

## 9 Author Contributions

Both authors designed the study, constructed the model, and provided the analysis. Both authors contributed to writing the manuscript. Both authors gave the final approval for publication.

## Supplementary Information

**Figure SI.1:**
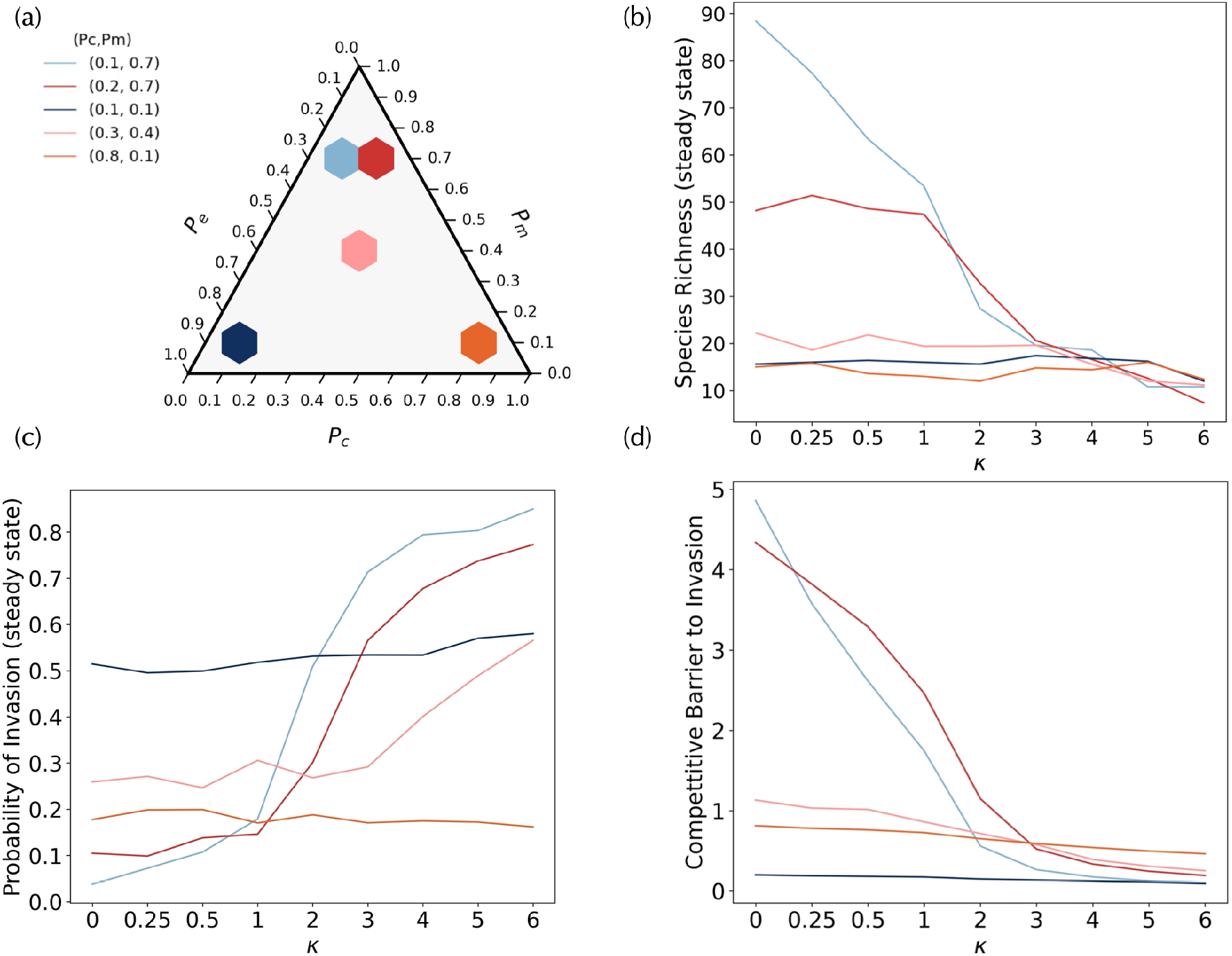
(a)Represents a ternary plot with selected microbial communities. The color of each selected community in the ternary plot corresponds to line plots in parts b-d, the line plots represent (b) species richness (at steady state), (c) invasion probability (at steady state), and (d) competitive barrier to invasion across a range of *κ* values.

**Figure SI.2:**
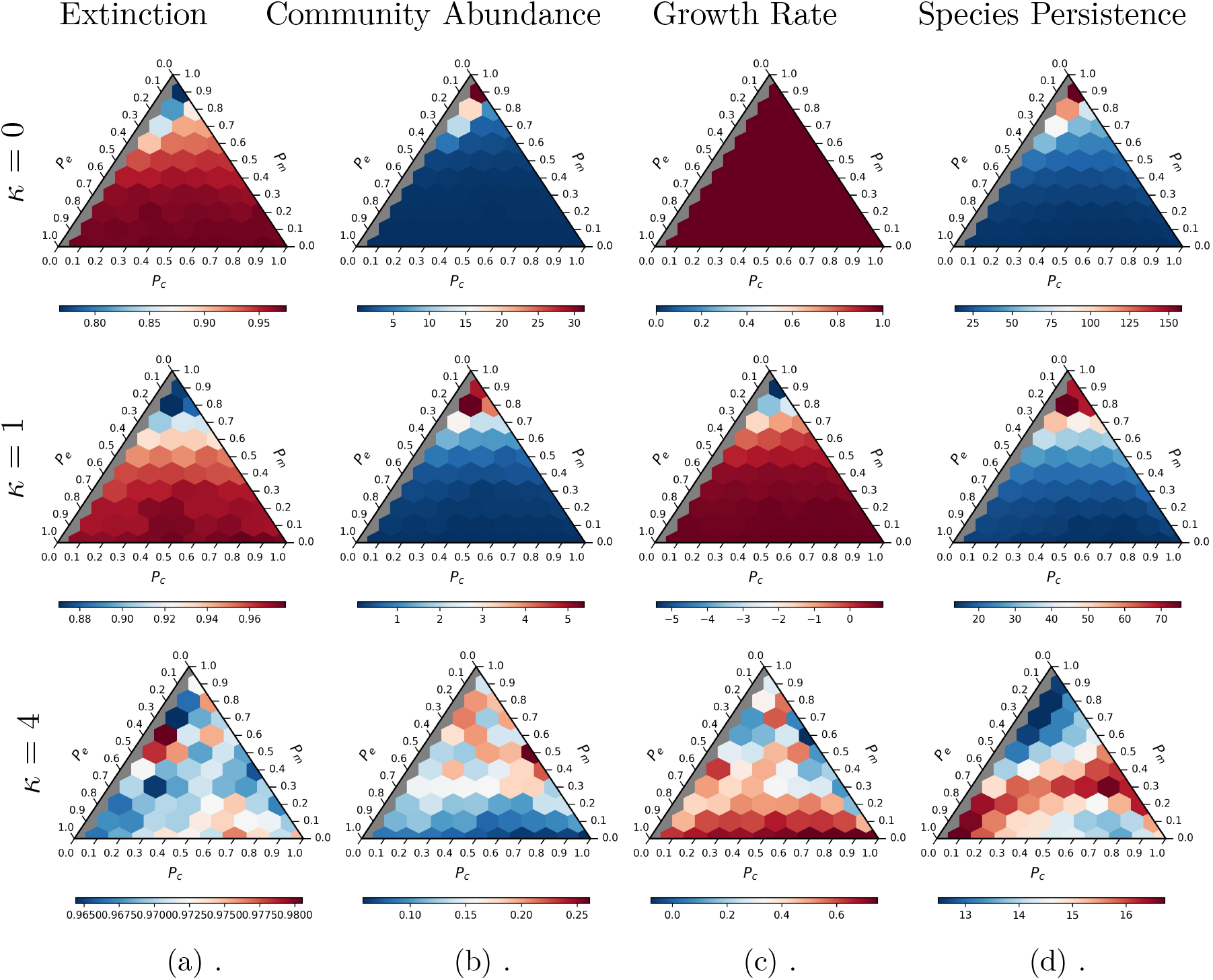
Ternary plots representing the host control on (a) probability of extinction (at steady state), (b) community abundance at steady state, and (c) growth rate, (d) mean species persistence for all possible communities with varying *P*_*c*_, *P*_*m*_ and *P*_*e*_. Note that scale of each plot shown above has a different range. Each row demonstrates an increasing host control strength (*κ*) for each community metric, from no host control (*κ* = 0) to high host control strength (*κ* = 4).

**Figure SI.3:**
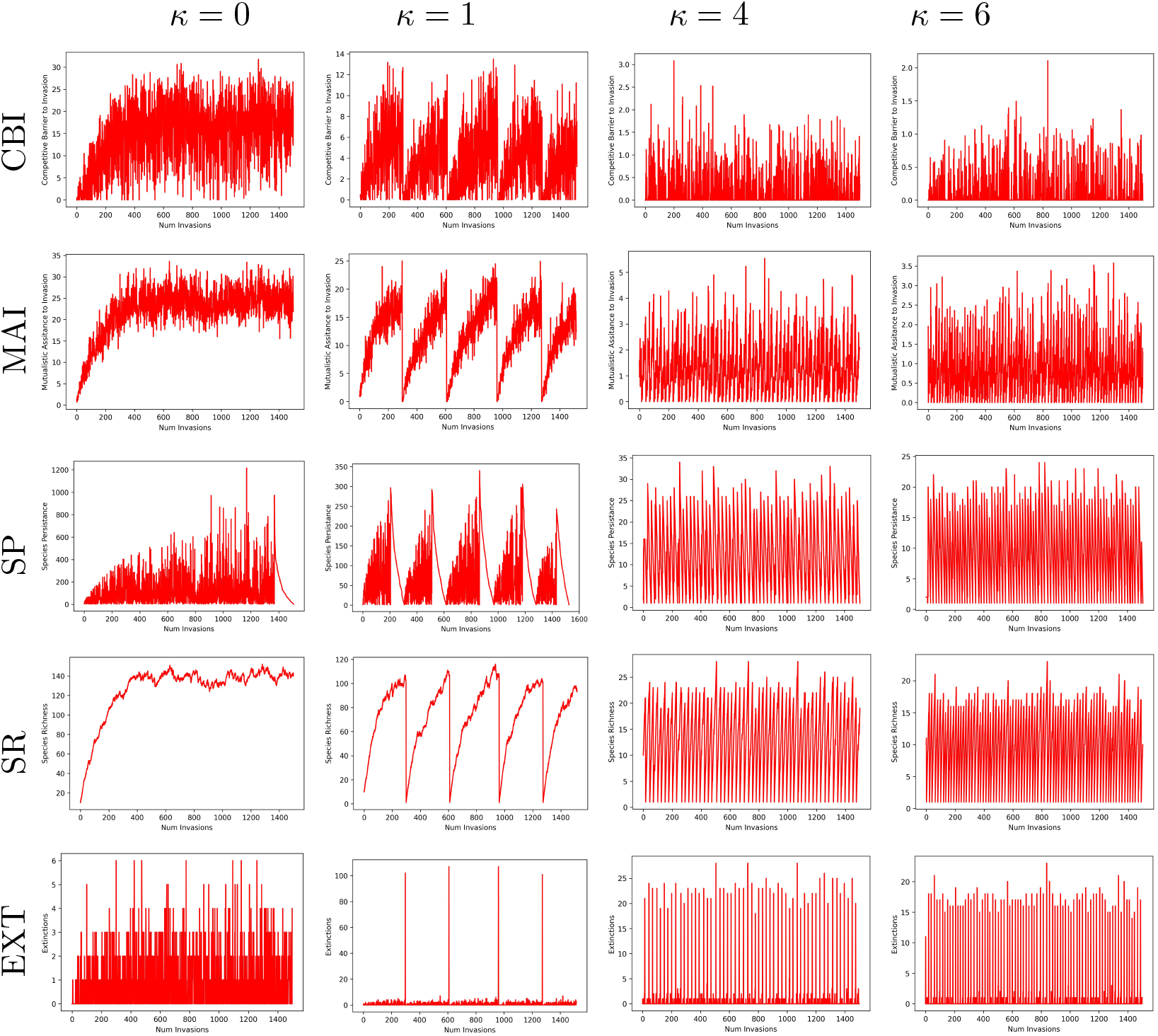
A series of line plots for the microbial community with *P*_*m*_ = 0.9 and *P*_*c*_ =0.1. The line plots represent how the community level properties, i) competitive barrier to invasion (CBI), ii) mutualistic assistance to invasion (MAI), iii) species persistence (SP), iv) species richness (SR) and v) extinction (EXT) vary across the sequential invasions in the community for a range of host control strength (*κ*) values.

### .1 Periodicity Analysis

We conducted periodicity analysis on the species richness of all microbial communities as a function of *κ*(0, 1, 4), where we determined the mode frequency and spectral density (amplitude) of species richness time series using the spectrum function in R. We observe frequent boom and bust cycles in species richness of highly mutualistic microbial communities, at high *κ* value.

**Figure SI.4:**
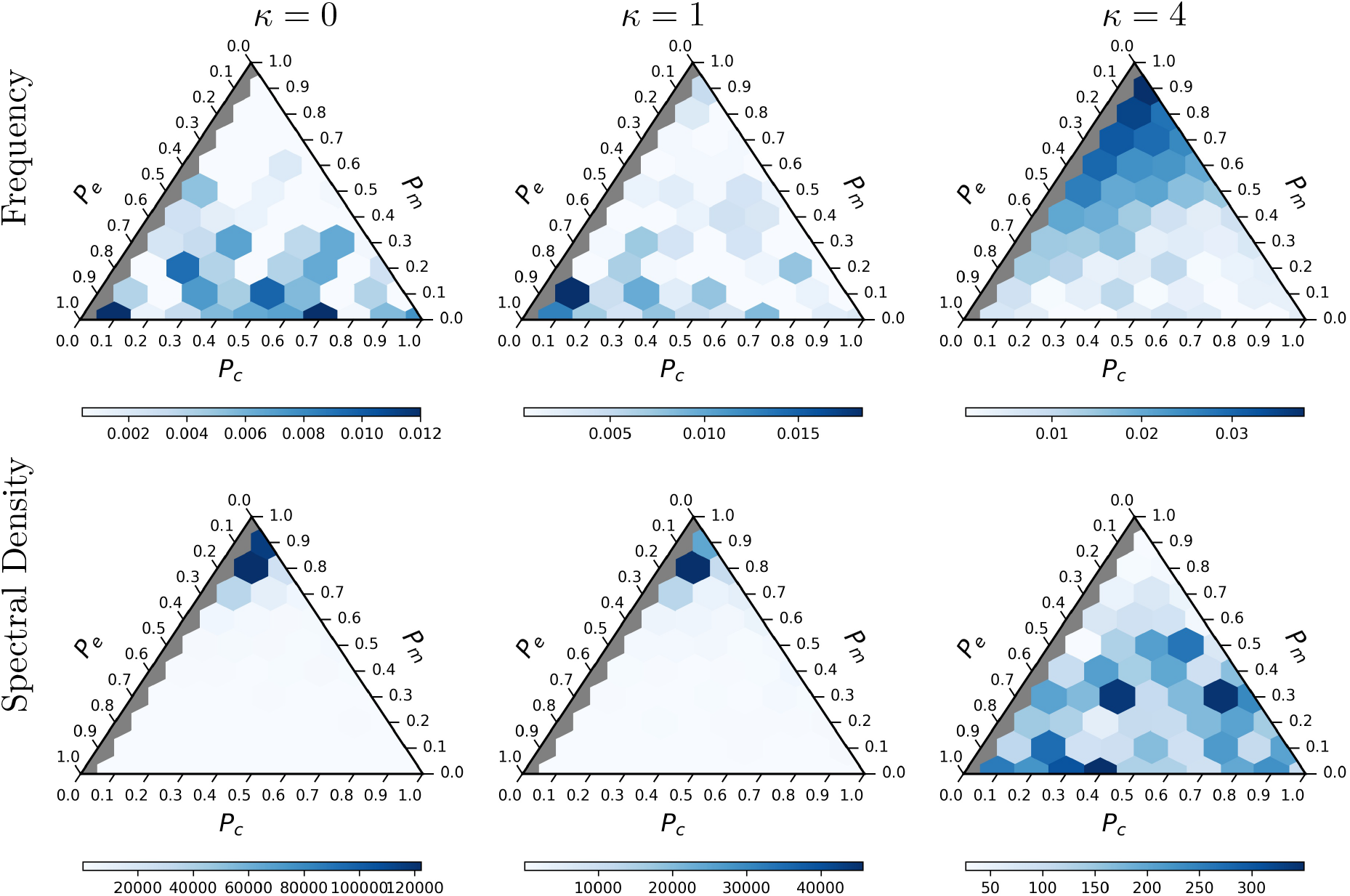
Ternary plot representing the frequency and spectral density for species richness time series observed for all possible microbial communities (with varying *P*_*c*_, *P*_*e*_, and *P*_*m*_) at different levels of host control strengths (*κ* = 0, 1, 4). Note that scale of each plot shown above has a different range.

### .1.1 Community Interactions under varying host control strength

We counted the number of interactions that were mutualistic, competitive and exploitative at the end of simulation for all communities at varying host control strength (*κ*).

**Figure SI.5:**
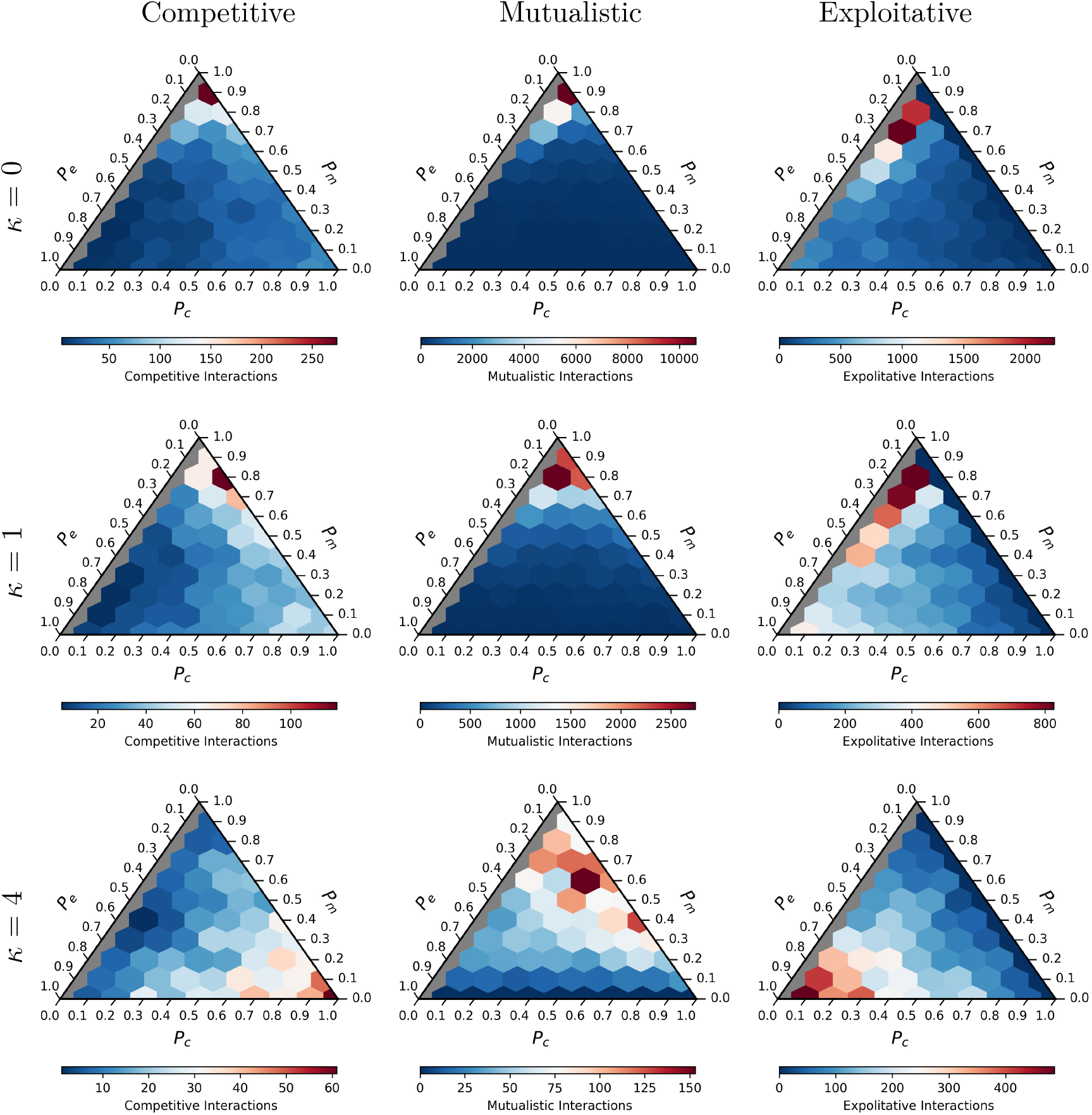
Number of competitive, mutualistic and exploitative interactions observed for each microbial community at the end of the community history under varying levels of host control (*κ* = 0, 1, 4). Note that scale of each plot shown above has a different range.

### .1.2 Changing host leash strength on the microbiome community

A healthy host can be perturbed to a disease like state which is known to shift microbiome community structure, size and diversity. Diseased-state is reflective of a stressed host, which can have an impaired host control on the microbiome. In our model, we varied the host leash (*κ*) to either a higher or a lower term (0, 1, 4) - encompassing possible variations in the host control, only after the initial community had reached a steady state. We chose the *κ* = 4, as the highest host control term, a value higher produces the same qualitative patterns. When changing the (*κ*), we allowed the community to reach equilibrium and then allowed sequential invasions to occur.

**Figure SI.6:**
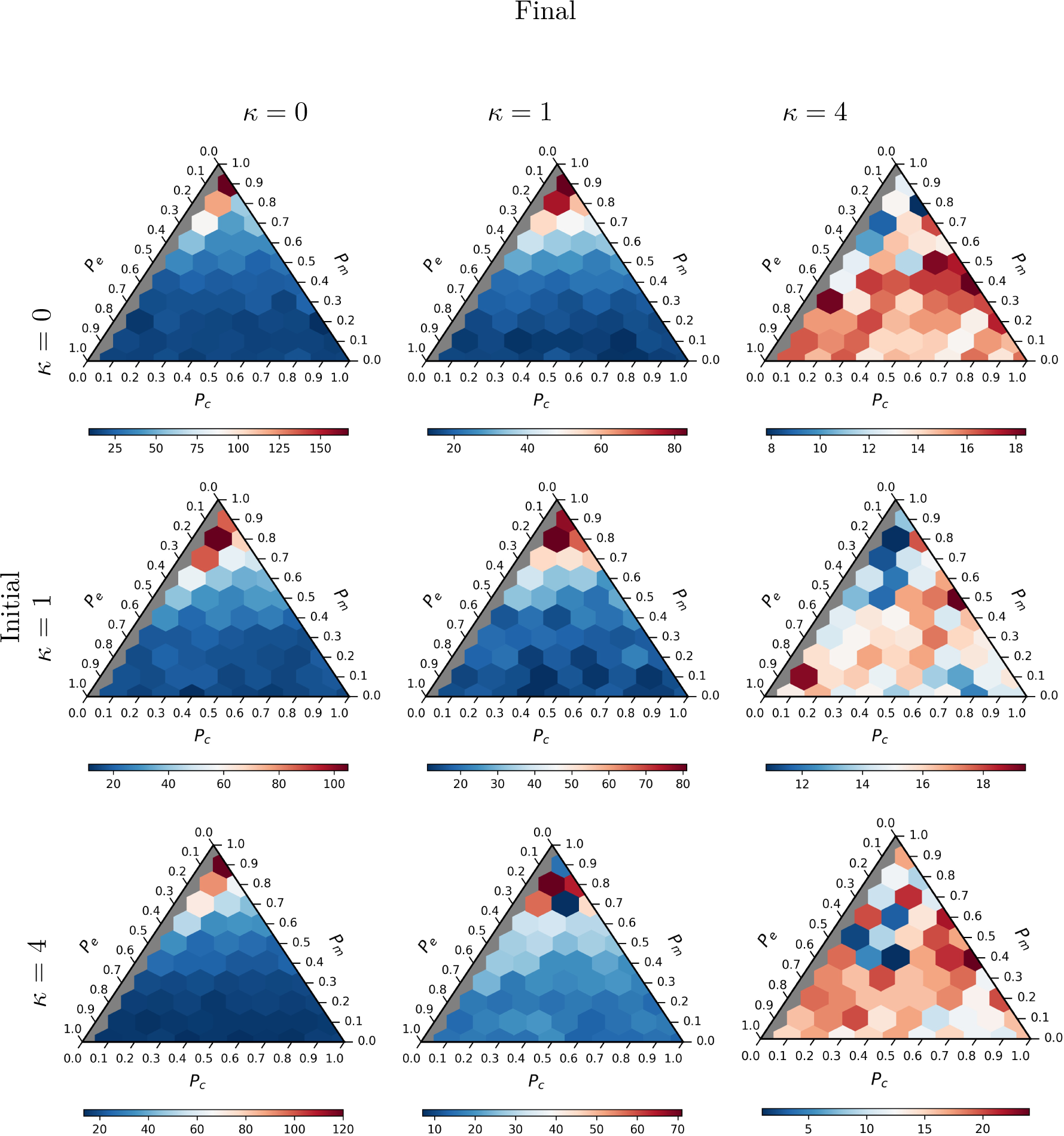
Ternary plots representing species richness (at steady state) observed for microbial communities. The *κ* terms observed on the vertical axis are the initial *κ* terms observed when the community reaches the steady state and the *κ* term on the horizontal axis represent the host control enforced on the community once after the the community has reached a steady state. Note that scale of each plot shown above has a different range.

**Figure SI.7:**
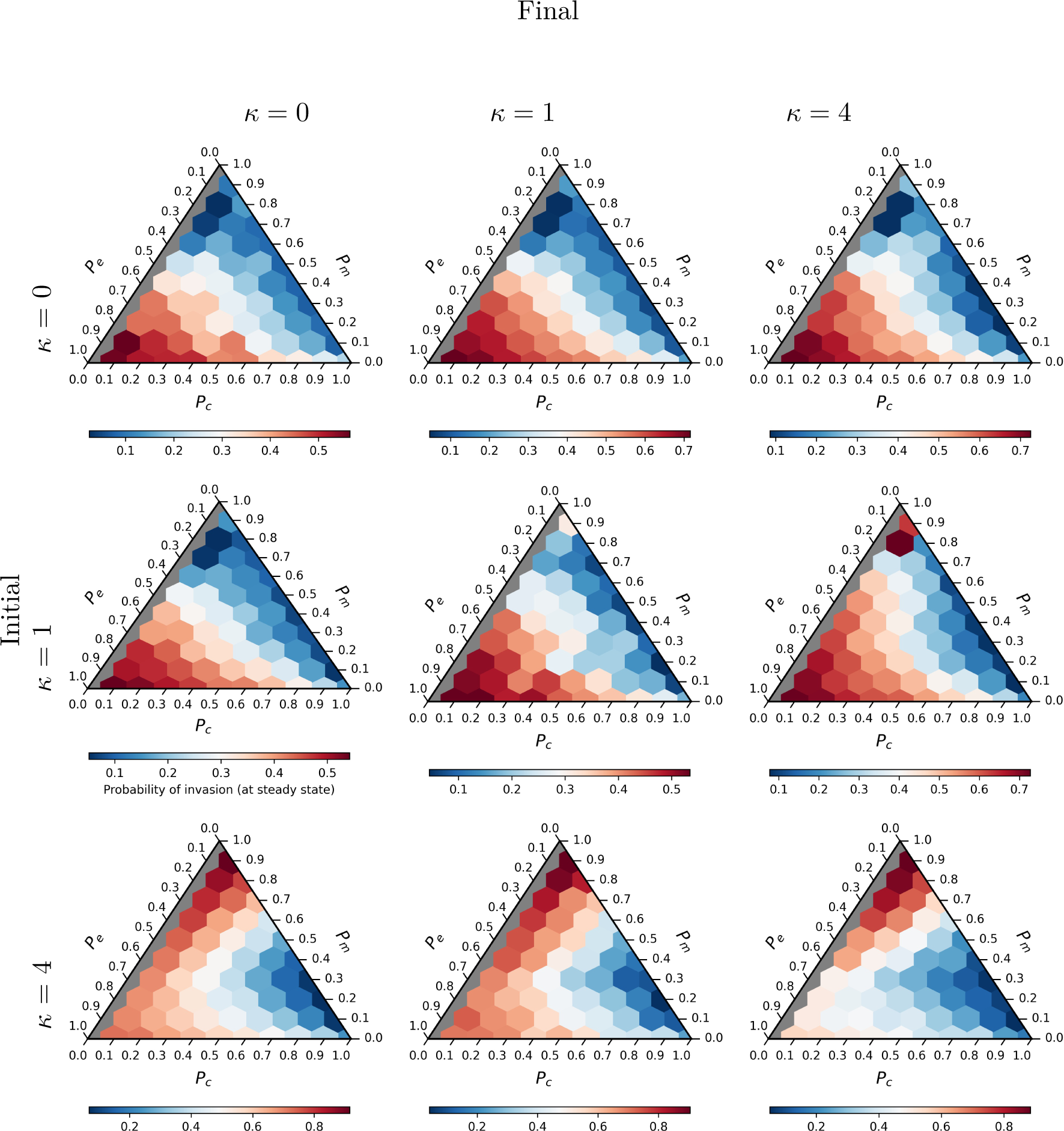
Ternary plots representing probability of invasion (at steady state) observed for microbial communities. The *κ* terms observed on the vertical axis are the initial *κ* terms observed when the community reaches the steady state and the *κ* term on the horizontal axis represent the host control enforced on the community once after the the community has reached a steady state. Note that scale of each plot shown above has a different range.

**Figure SI.8:**
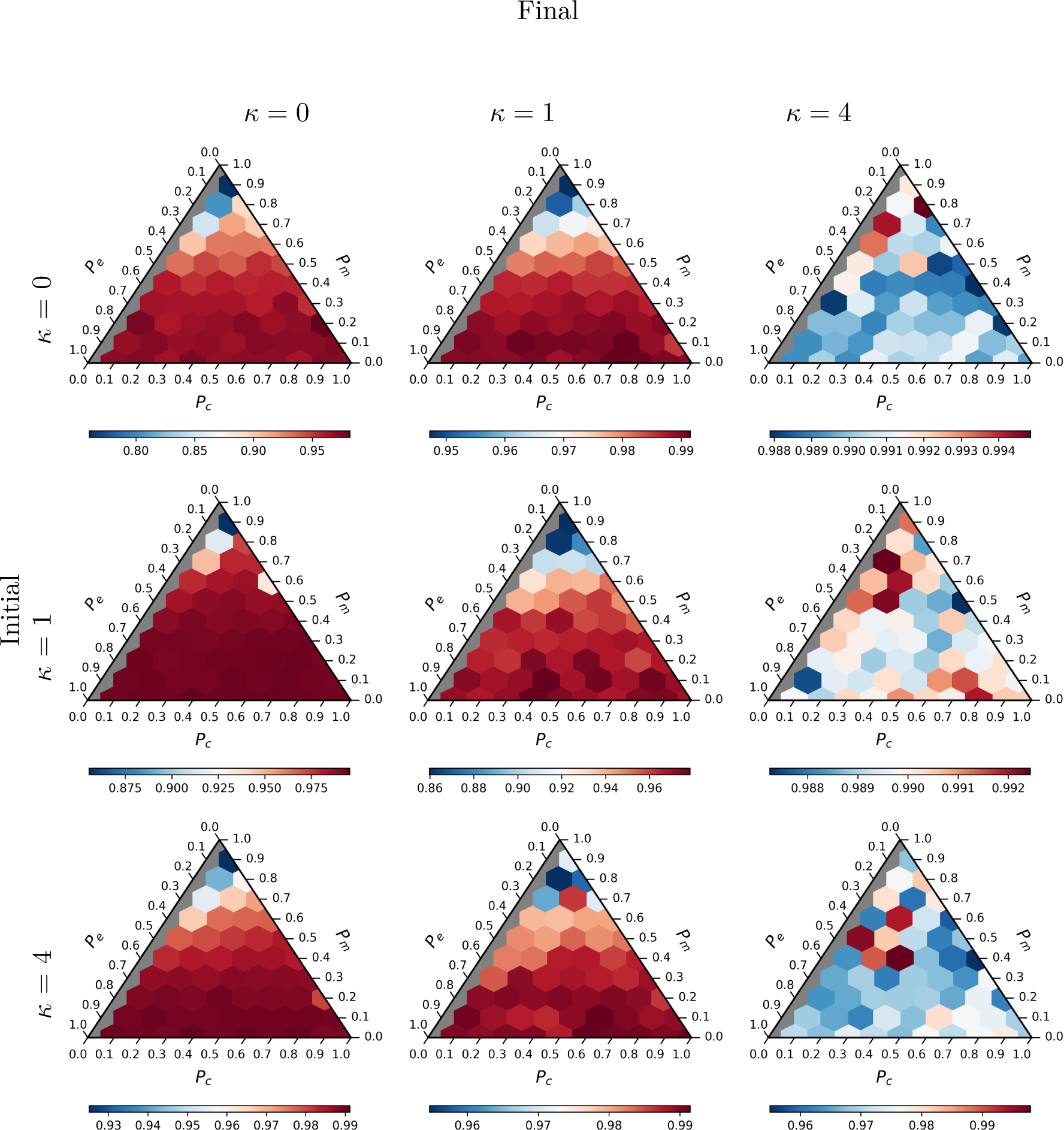
Ternary plots representing probability of extinction (at steady state) observed for microbial communities.The *κ* terms observed on the vertical axis are the initial *κ* terms observed when the community reaches the steady state and the *κ* term on the horizontal axis represent the host control enforced on the community once after the the community has reached a steady state. Note that scale of each plot shown above has a different range.

**Figure SI.9:**
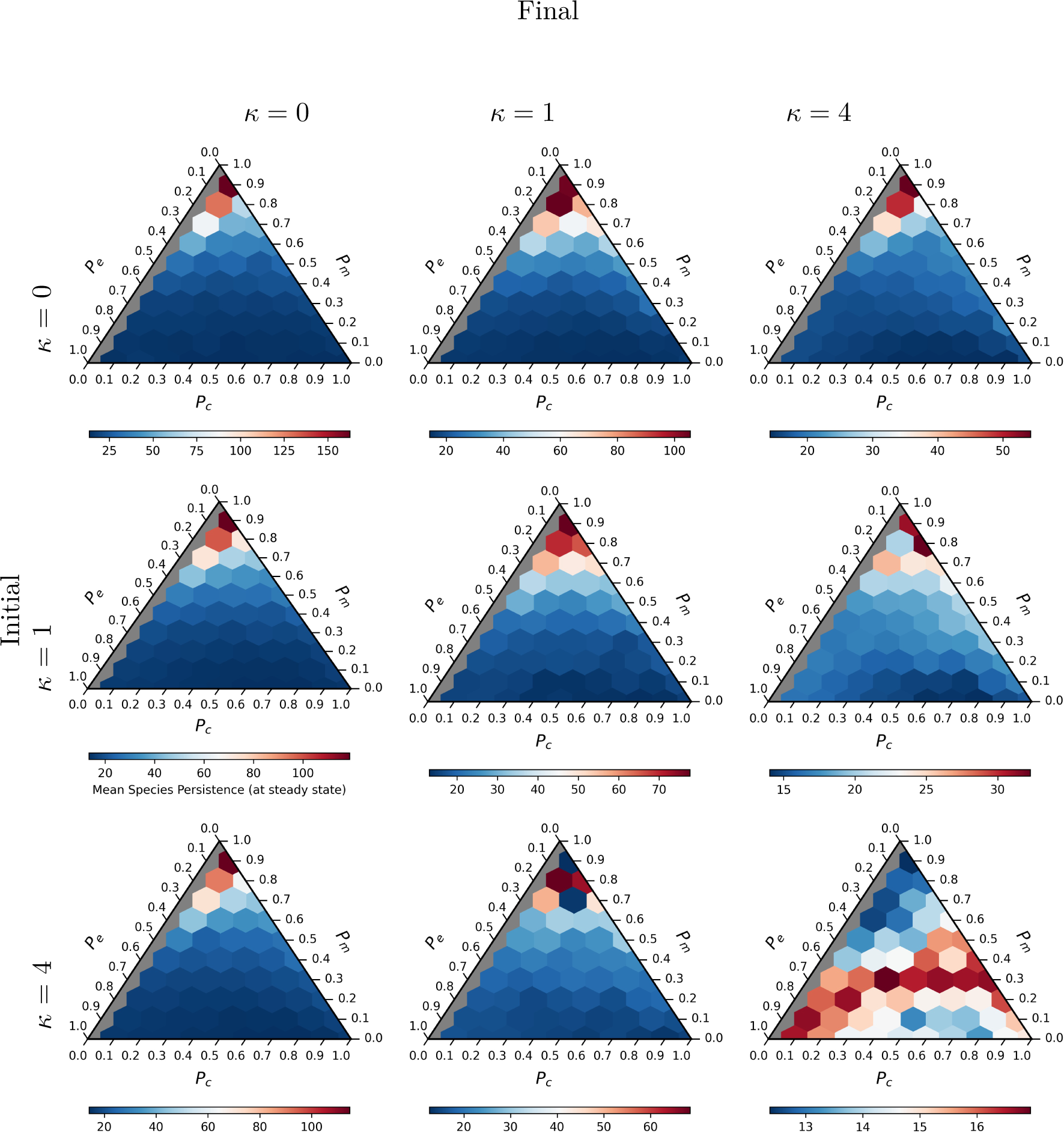
Ternary plots representing mean species persistence (at steady state) observed for microbial communities. The *κ* terms observed on the vertical axis are the initial *κ* terms observed when the community reaches the steady state and the *κ* term on the horizontal axis represent the host control enforced on the community once after the the community has reached a steady state. Note that scale of each plot shown above has a different range.

### .1.3 Multi-stability Analysis

**Figure SI.10:**
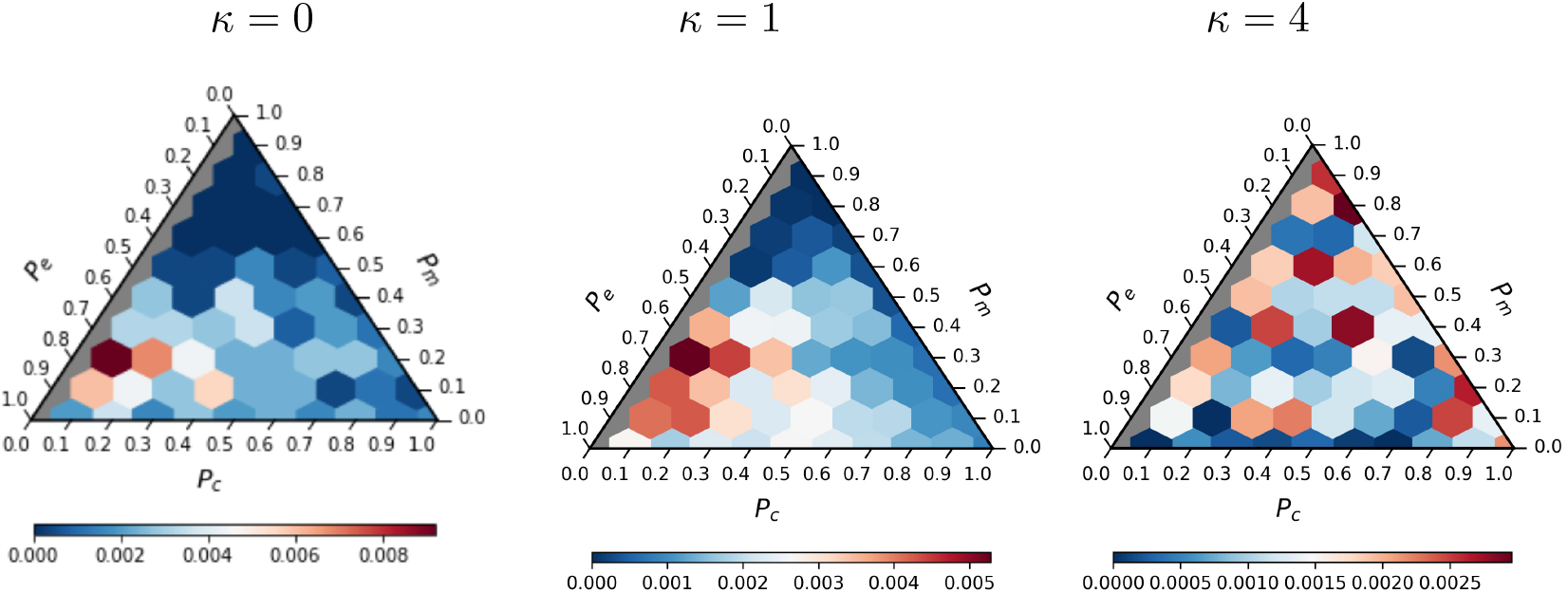
Represents ternary plots showing the probability that an equilibrium in the community history shows oscillations in population dynamics for all microbial communities across a range of *κ* values..

**Figure SI.11:**
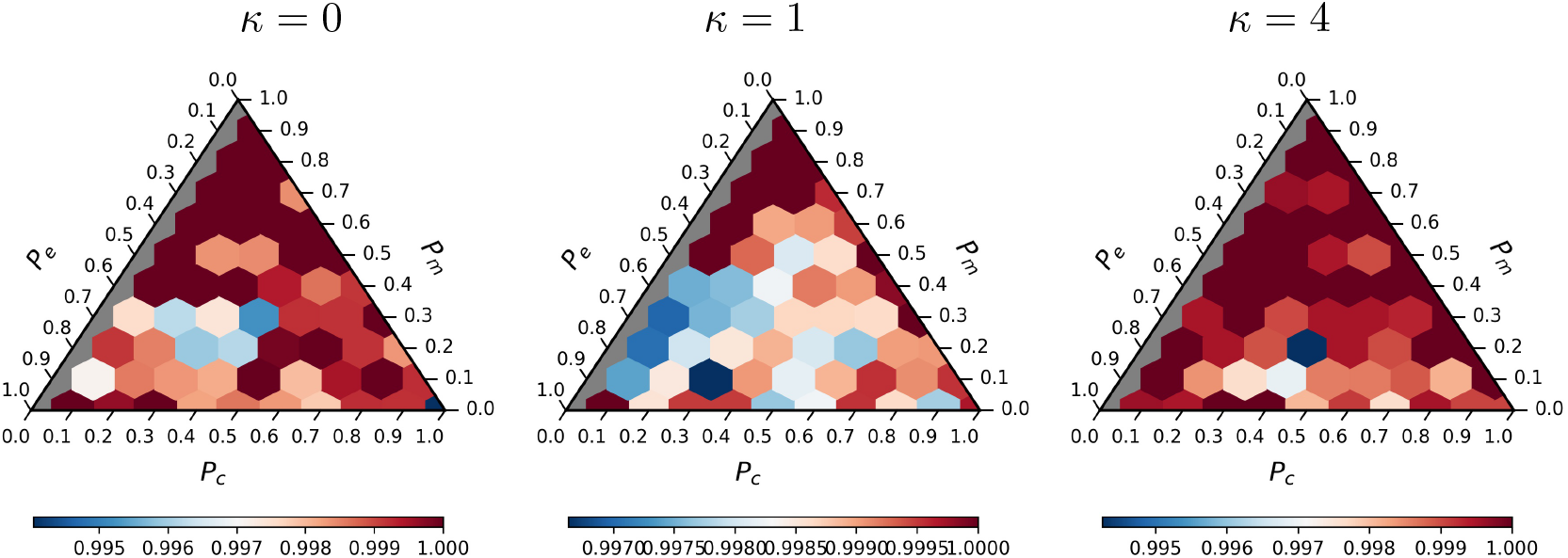
Represents ternary plots showing the ratio of stable equilibria observed for microbial communities when assayed with 25 different initial population sizes across a range of *κ* values.

